# GABAergic neurons in the olfactory cortex projecting to the lateral hypothalamus in mice

**DOI:** 10.1101/413328

**Authors:** Koshi Murata, Tomoki Kinoshita, Yugo Fukazawa, Kenta Kobayashi, Kazuto Kobayashi, Kazunari Miyamichi, Hiroyuki Okuno, Haruhiko Bito, Yoshio Sakurai, Masahiro Yamaguchi, Kensaku Mori, Hiroyuki Manabe

## Abstract

Olfaction guides goal-directed behaviours including feeding. To investigate how central olfactory neural circuits control feeding behaviour in mice, we performed retrograde tracing from the lateral hypothalamus (LH), an important feeding centre. We observed a cluster of retrogradely labelled cells distributed in the posteroventral region of the olfactory peduncle. Histochemical analyses revealed that a majority of these retrogradely labelled projection neurons expressed glutamic acid decarboxylase 65/67 (GAD65/67), but not vesicular glutamate transporter 1 (VGluT1). We named this region with GABAergic projection neurons the ventral olfactory nucleus (VON) to discriminate it from the conventional olfactory peduncle. VON neurons were less immunoreactive for DARPP-32, a striatal neuron marker, in comparison to those in the olfactory tubercle and nucleus accumbens, which distinguished the VON from the ventral striatum. Fluorescent labelling confirmed synaptic contacts between VON neurons and olfactory bulb projection neurons. Rabies-virus-mediated trans-synaptic labelling revealed that VON neurons received synaptic inputs from the olfactory bulb, other olfactory cortices, horizontal limb of the diagonal band, and prefrontal cortex. Collectively, these results identified novel GABAergic projection neurons in the olfactory cortex that can integrate olfactory sensory and top-down inputs and send inhibitory output to the LH, which may contribute to forming odour-guided LH-related behaviours.

## Introduction

The central olfactory system translates odour information into motivated behaviours, including appetite-based food approach and eating behaviours^1^. Recent studies have revealed neuronal circuit mechanisms by which odorants evoke specific behaviours, such as fear responses to predator odours^2,3^ and attractive responses to social odours^4^. However, it is still unclear how the central olfactory neural circuits control feeding-related behaviours in mammals.

Odorants activate olfactory sensory neurons and coded by activation of specific combinations of glomeruli in the olfactory bulb (OB), the first relay centre of the central olfactory system^5^. Mitral cells and tufted cells (M/TCs) are projection neurons in the OB. They convey odour information to several areas in the olfactory cortex which is composed of the anterior olfactory nucleus (AON), tenia tecta, dorsal peduncular cortex, anterior piriform cortex (APC), olfactory tubercle (OT), posterior piriform cortex (PPC), cortical amygdaloid nuclei, and entorhinal cortex^6^.

We previously reported that c-fos activity increased in the anteromedial domain of the OT when mice showed attractive responses to a learned odour cue that predicted food^7^. The OT has indirect anatomical connections to the lateral hypothalamus (LH), an important feeding centre^8,9^, via the ventral pallidum^10^. Knowledge of neural pathways from the olfactory cortex to the LH is crucial to identify how olfactory information is translated into feeding-related behaviours. Price et al. examined the neural connections between the central olfactory system and the LH in rats^11^, and reported that several parts of the olfactory cortex have axonal projections to the LH. Since then, however, no neuroanatomical studies have addressed how olfactory information is conveyed to the LH in mammals.

Here, we re-examined the neural pathways from the central olfactory system to the LH in mice using cholera toxin B subunit (CTB), and viral and genetic techniques to trace neural circuits. In agreement with the previous study^11^, we observed a subset of retrogradely labelled cells clustered in a postero-ventral region of the olfactory peduncle. Our analyses revealed that a majority of LH-projecting neurons in this region were GABAergic neurons, in sharp contrast to the AON and ventral tenia tecta (VTT) where principal neurons are glutamatergic. We also found that the GABAergic neurons extended dendrites to layer I and received synaptic inputs from the olfactory bulb neurons. These results suggest a novel population of GABAergic neurons in the olfactory cortex projecting to the LH.

## Results

### Retrograde tracer injection into the LH revealed a cluster of GABAergic neurons in the olfactory peduncle

To reveal neural pathways from the olfactory cortex to the LH, we injected a retrograde tracer, CTB conjugated with Alexa 555, into the mouse LH. We targeted an area of the LH where orexin neurons and melanin-concentrating hormone (MCH) neurons were distributed (Fig. 1A), as both these neuronal subpopulations are involved in feeding behaviours^12,13^. CTB-labelled cells were widely distributed in the brain including the prefrontal cortex (Fig. 1B, left). Guided by previous studies reporting that the olfactory cortex has axonal projections to the LH^11^, we found a cluster of CTB-labelled cells in an area surrounded by the AON, APC, VTT, and OT (Fig. 1B right). The size of CTB-labelled somata was smaller than that of principal neurons in the posterior part of the AON (Figs. 1B and 2), suggesting that their neuronal subtypes differed from principal neurons in the AON.

**Figure 1.**
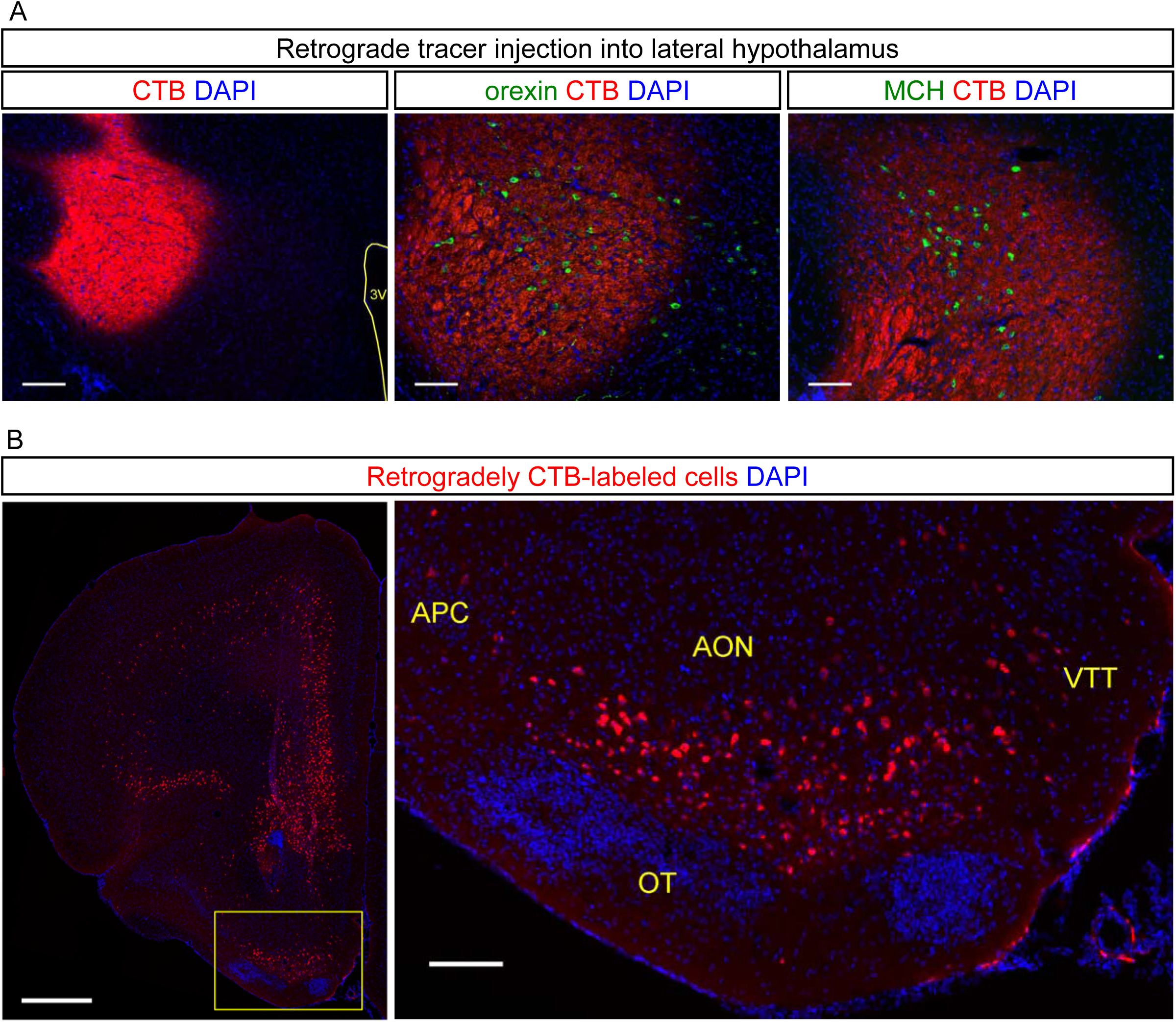
Retrograde tracer revealed a cluster of neurons projecting to the lateral hypothalamus (LH) surrounded by the olfactory cortex. (A) Coronal sections of the LH after injection of Alexa 555-conjugated cholera toxin B subunit (CTB, red) with DAPI staining (blue). Immunostaining for orexin (middle, green) or melanin-concentrating hormone (MCH) (right, green) was performed. Scale bars: 200 μm in left panel, 100 μm in middle and right panels. 3V, third ventricle. (B) CTB-labelled cells in the frontal cortex and olfactory cortex. The inset in the left panel is magnified in the right panel. APC, anterior piriform cortex; AON, anterior olfactory nucleus; VTT, ventral tenia tecta; OT, olfactory tubercle. Scale bars: 500 μm in left panel, 100 μm in right panel.

To examine whether the CTB-labelled cells were glutaminergic or GABAergic, we performed *in situ* hybridization for mRNA of vesicular glutamate transporters (VGluTs) and glutamic acid decarboxylase (GAD) 65/67 in this area (Fig. 2). VGluT1-expressing cells were distributed in the dorsal edge of the area in which CTB-labelled cells were clustered. We observed a cluster of GAD65/67-expressing cells just above the rostral tip of the OT, which overlapped with the distribution of CTB-labelled cells. VGluT2-and VGluT3-expressing cells were scarcely observed in this area. We then directly examined expression of VGluT1 and GAD65/67 mRNAs in CTB-labelled cells by double fluorescent labelling (Fig. 2B). VGluT1-expressing CTB-labelled cells were distributed in the ventral border of the AON, and 7.8% of the CTB-labelled cells were VGluT1-positive (n = 154 cells from two mice). In contrast, 84.6% of the CTB-labelled cells were GAD65/67-positive (n = 214 cells from three mice), which suggested that principal neurons in this area were GABAergic. Projection neurons in the olfactory cortex, except for the OT, are thought to be glutamatergic. This cellular profile raised a possibility that the GABAergic CTB-labelled cells did not belong to the conventional olfactory peduncle. We therefore temporarily named this area the ventral olfactory nucleus (VON), because the cluster of GABAergic neurons projecting to the LH was located just ventral to the AON.

**Figure 2.**
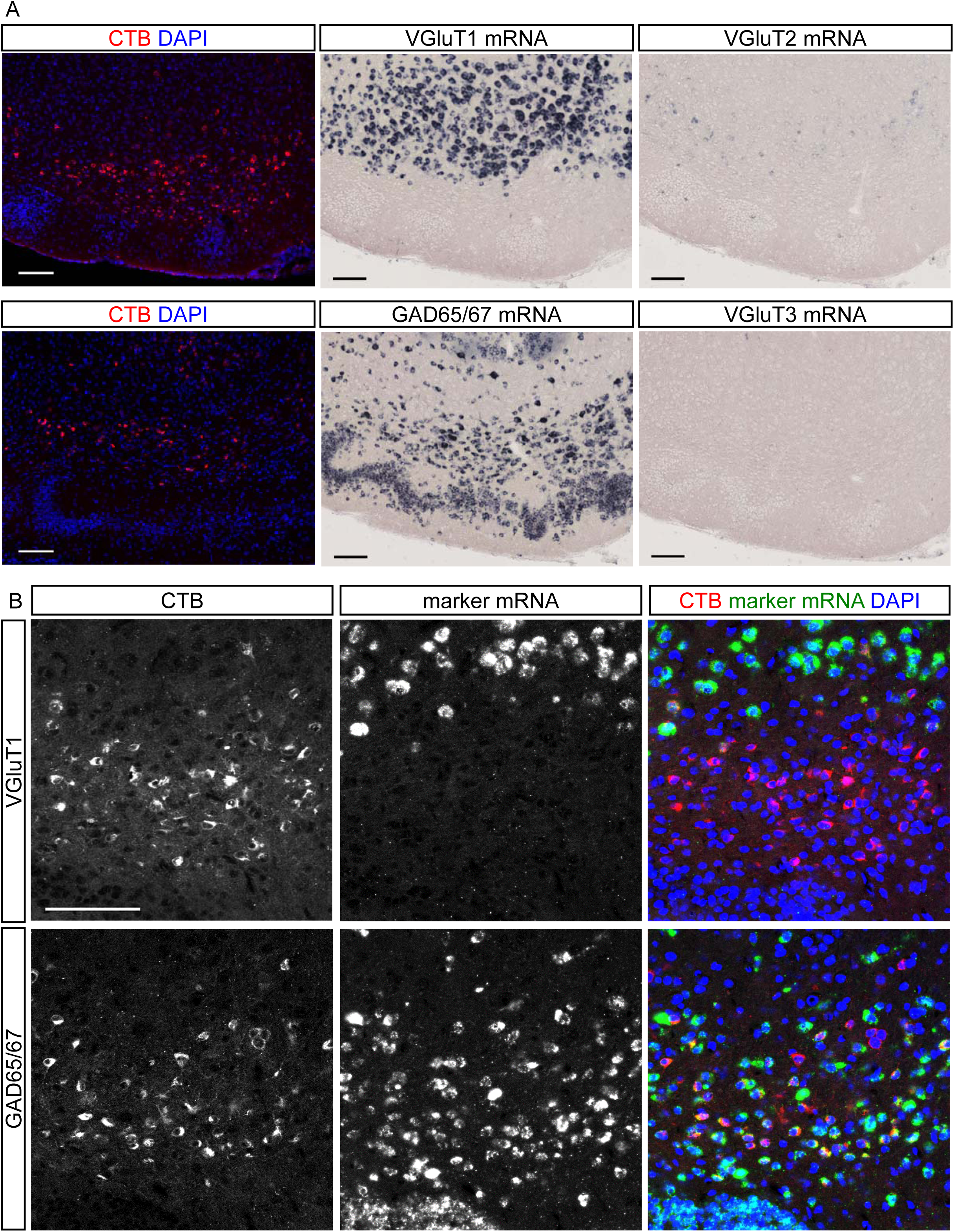
Majority of CTB-labelled neurons in the ventral olfactory nucleus (VON) were GABAergic. (A) *In situ* hybridization for VGluTs and GAD65/67 mRNAs, and adjacent slice images of cholera toxin B subunit (CTB)-labelled ventral olfactory nucleus (VON) neurons. Scale bars: 100 μm. (B) Double fluorescent labelling of CTB and VGluT1 (upper panels) or GAD65/67 (lower panels) mRNAs with DAPI staining. Scale bar: 100 μm.

One possible cellular profile of GABAergic neurons in the VON is medium spiny neurons in the OT and nucleus accumbens (NAc), namely the ventral striatum^14,15^. To examine whether the CTB-labelled cells in the VON were different from medium spiny neurons of the ventral striatum, we performed immunostaining for DARPP-32, a marker for striatal neurons^16^, alongside CTB labelling (Fig. 3). CTB-labelled cells were distributed in both DARPP-32 strongly immunoreactive and less immunoreactive regions, which corresponded to the ventral striatum and VON, respectively. These results support the idea that GABAergic neurons in the VON formed distinct cellular populations from medium spiny neurons in the OT and NAc.

**Figure 3.**
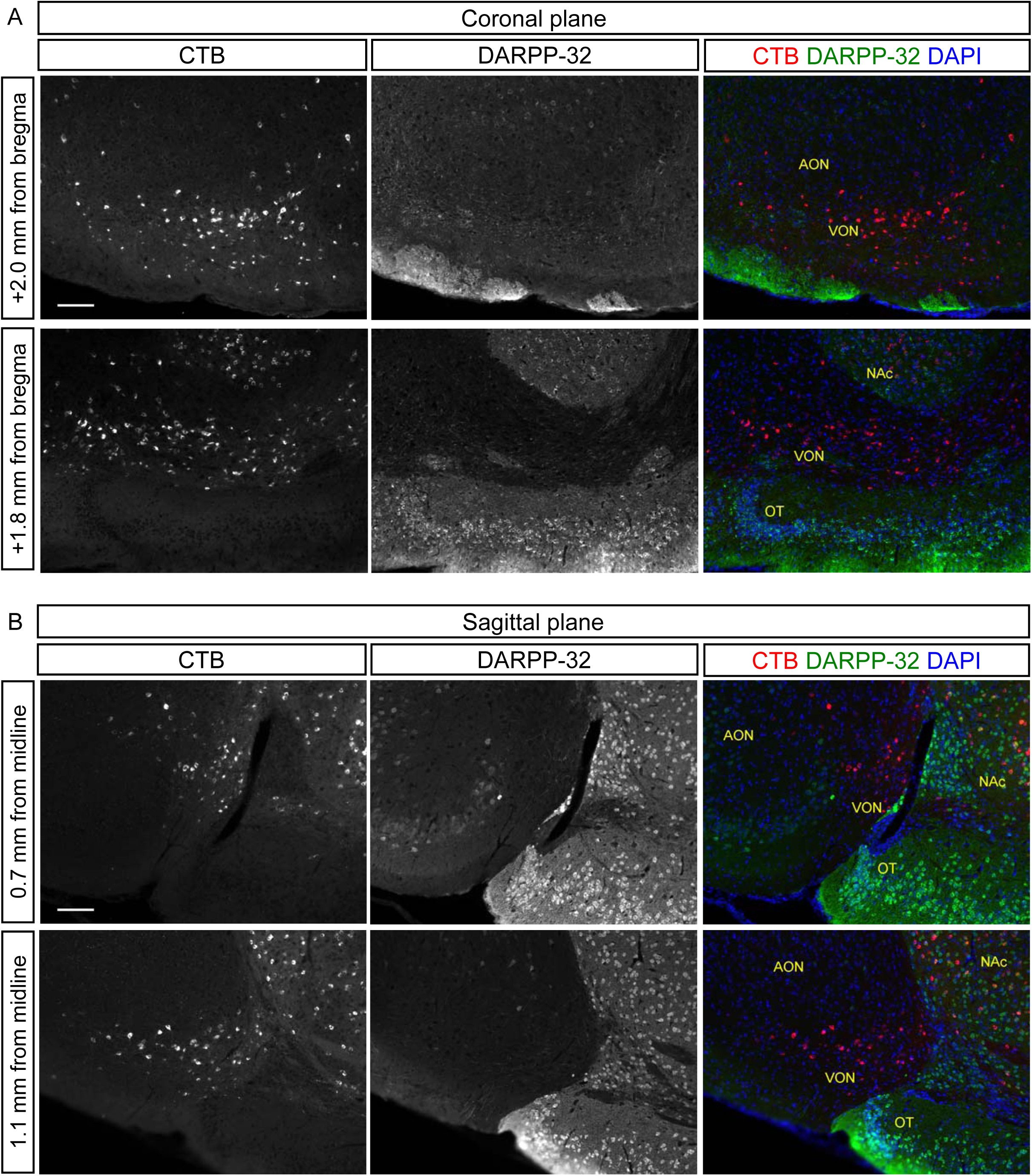
The VON was distinguishable from the ventral striatum by DARPP-32 immunoreactivity. (A and B) Coronal sections (A) and sagittal sections (B) of the VON after injection of Alexa 555-conjugated cholera toxin B subunit (CTB) (red) into the lateral hypothalamus (LH) with immunostaining for DARPP-32 (green) and DAPI staining (blue). AON, anterior olfactory nucleus; VON, ventral olfactory nucleus; OT, olfactory tubercle; NAc, nucleus accumbens. Scale bars: 100 μm.

### Synaptic contacts from OB M/TCs to VON neurons

Olfactory cortex is defined as those areas that receive direct synaptic inputs from M/TCs, projection neurons in the OB^6^. To reveal synaptic contacts from M/TCs onto the dendrites of VON neurons, we used a transgenic mouse line in which M/TCs were labelled by tdTomato (Pcdh21-nCre; tdTomato Cre reporter line, Ai14^17,18^). We injected retrogradely-spreading adeno-associated virus (AAV) encoding EGFP (AAVrg-CAG-EGFP^19^) into the LH of the transgenic mice (Figs. 4-5). EGFP-labelled dendrites of the VON neurons were innervated by tdTomato-labelled axons of M/TCs (Fig. 4A, B), suggesting that VON neurons receive axonal synaptic inputs in layer Ia of the olfactory cortex. Notably, dendrites of VON neurons did not innervate DARPP-32 strongly immunoreactive regions, the NAc and OT (Fig. 4B).

**Figure 4.**
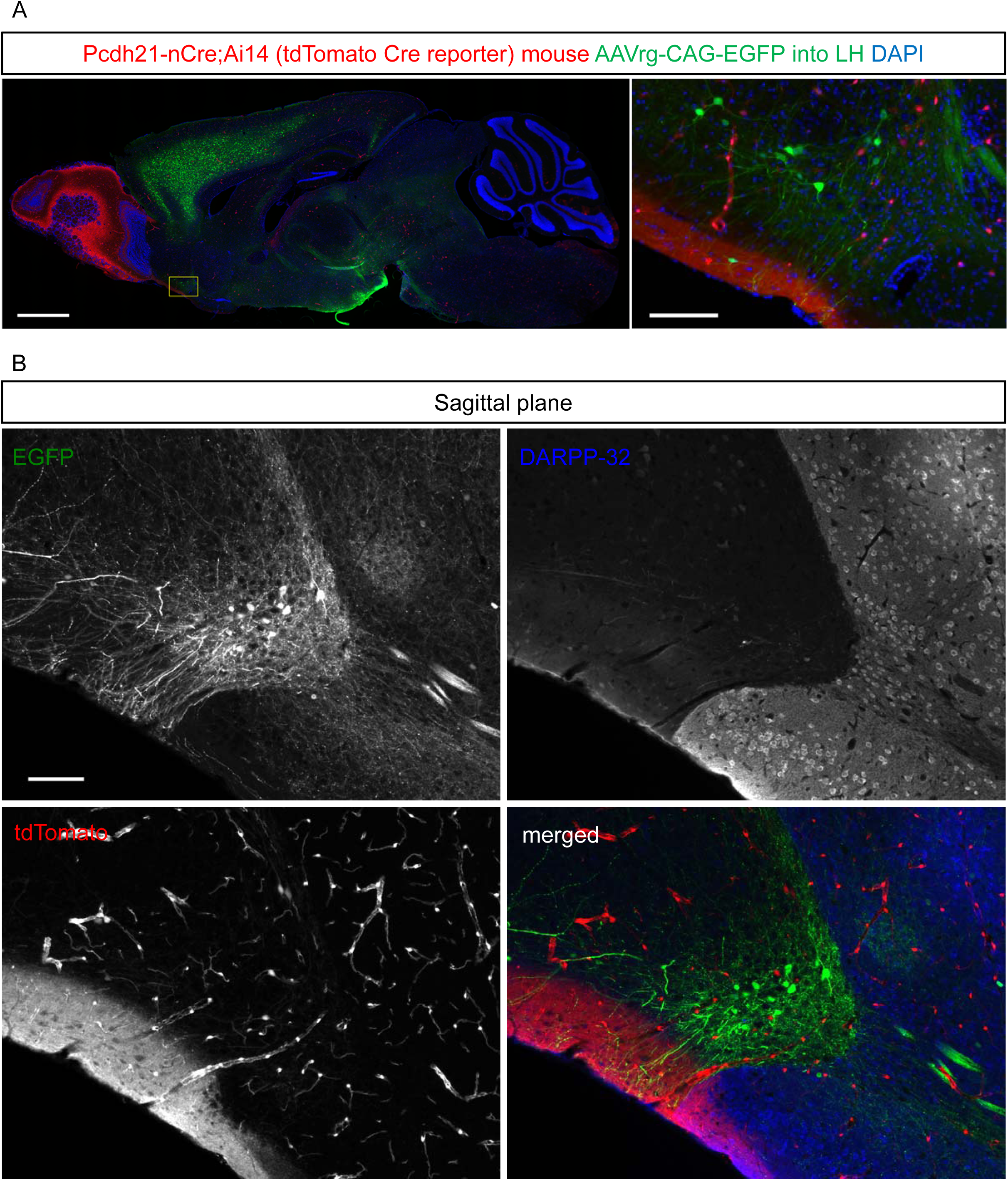
Mitral and tufted cells in the olfactory bulb send axons to the VON. (A) Sagittal sections of the whole brain of the mitral and tufted cells (M/TCs) in a tdTomato-labelled transgenic mouse after injection of retrograde adeno-associated virus (AAV) vector encoding EGFP into the lateral hypothalamus (LH). The inset of the left panel is magnified in the right panel. We note that vascular endothelial cells and some neurons in the olfactory cortex were labelled by tdTomato as well as M/TCs. Scale bars: 1 mm in left panel, 100 μm in right panel. (B) Sagittal sections of the ventral olfactory nucleus (VON) with immunostaining for DARPP-32 (blue). Dendrites of EGFP-labelled VON neurons (green) innervated layer Ia of the olfactory cortex which was innervated by axons of tdTomato-labelled M/TCs (red). Scale bar: 100 μm.

**Figure 5.**
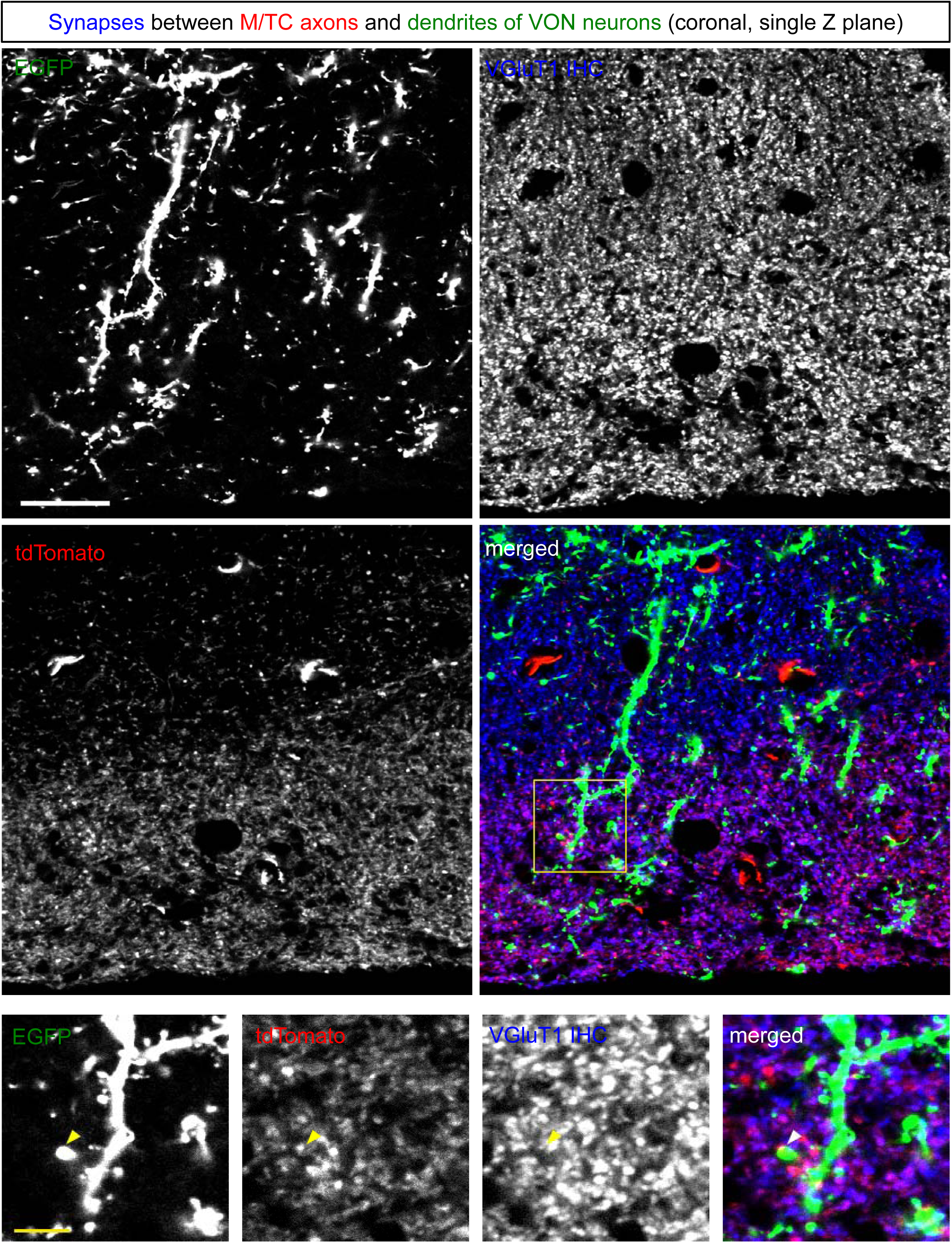
Synaptic contacts between mitral and tufted cells and spines of neurons in the VON. Single place confocal images of dendrites of EGFP-labelled ventral olfactory nucleus (VON) neurons (green), axons of tdTomato-labelled mitral and tufted cells (M/TCs) (red) and immunoreactivity for vesicular glutamate transporter 1 (VGluT1) (blue) in layer Ia of the VON. Lower four panels are magnified views of the inset. Scale bars: 20 μm for upper four panels, 5 μm for lower four panels.

We next performed immunostaining for VGluT1, a presynaptic marker of M/TCs^20,21^, and examined apposition of VGluT1 signals and dendrites of VON neurons (Fig. 5). EGFP-labelled dendrites of VON neurons had spines (Figs. 4B and 5). Axonal boutons of tdTomato-labelled M/TCs in layer Ia were apposed to dendritic spines of VON neurons, which colocalised with VGluT1 immunoreactivity (arrowheads in Fig. 5, lower panels). These results suggest that VON neurons receive glutamatergic inputs from M/TCs, and support the idea that the VON is a part of the olfactory cortex.

### Trans-synaptic retrograde tracing from the VON using a modified rabies virus

To further confirm whether the VON was a part of the olfactory cortex, namely whether VON neurons received direct synaptic inputs from M/TCs in the OB, we performed trans-synaptic labelling of VON neurons combining an EnvA-pseudotyped glycoprotein-deleted rabies virus encoding EGFP (SAD-dG-EGFP+EnvA) with AAV-mediated expression of the EnvA receptor (TVA) and rabies glycoprotein^22^-^25^. This labelling also revealed input pathways to the VON neurons in whole brain regions. To achieve selective initial infection of VON neurons by the rabies virus, we used the neural pathway specific-tracing method (tracing the relationship between input and output, TRIO^26^). We first injected a retrograde Cre-coding lentiviral vector (NeuRet-Cre^27^) into the LH, and Cre-dependent AAV vectors encoding TVA-mCherry and rabies G into the VON. Two weeks after the injection, we injected the modified rabies virus (SAD-dG-EGFP+EnvA) into the VON (Fig. 6A).

**Figure 6.**
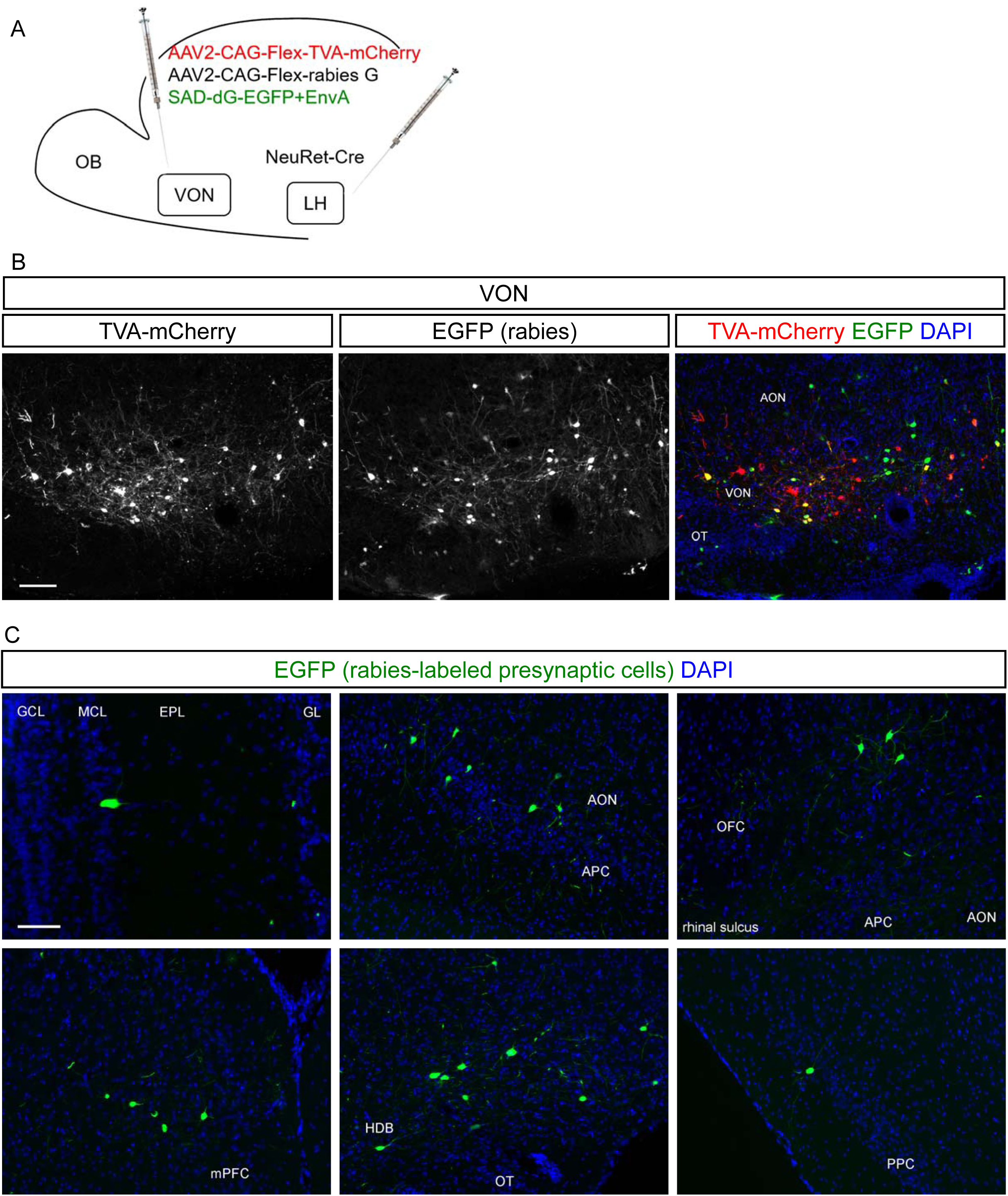
Rabies virus-mediated trans-synaptic retrograde tracing from the VON. (A) Schema of virus-mediated neuronal pathway-specific retrograde tracing. We first injected a retrograde lentiviral vector coding Cre (NeuRet-Cre) into the lateral hypothalamus (LH), and Cre-dependent adeno-associated virus (AAV) vector TVA-mCherry and rabies G into the VON. Two weeks after the first injection, EnvA-pseudotyped glycoprotein-deleted rabies virus encoding EGFP was injected into the VON. We then observed distribution of EGFP-labelled presynaptic cells throughout the whole brain. (B) Coronal sections of the VON. Left, TVA-mCherry expressing neurons; middle, rabies-derived EGFP expressing neurons; right, colour merged with DAPI staining. Yellow cells are TVA-mCherry(+) EGFP(+) starter cells. AON, anterior olfactory nucleus; VON, ventral olfactory nucleus; OT, olfactory tubercle. Scale bar: 100 μm. (C) Distribution of EGFP-labelled presynaptic cells to the VON. GCL, granule cell layer; MCL, mitral cell layer; EPL, external plexiform layer; GL, glomerular layer; AON, anterior olfactory nucleus; APC, anterior piriform cortex; OFC, orbitofrontal cortex; mPFC, medial prefrontal cortex; HDB, horizontal limb of the diagonal band; OT, olfactory tubercle; PPC, posterior piriform cortex. Scale bar: 100 μm.

Starter cells of TVA-mCherry(+) and EGFP(+) cells were observed in the VON (Fig. 6B). We detected EGFP-labelled cells in the OB (Fig. 6C, upper left panel), confirming that the VON neurons receive direct synaptic input from the OB. EGFP-labelled cells were also distributed in the AON and OT (Fig. 6B, right panel), APC (Fig. 6C, upper middle panel), orbitofrontal cortex (Fig. 6C, upper right panel), medial prefrontal cortex (Fig. 6C, lower left panel), horizontal limb of the diagonal band (Fig. 6C, lower middle panel), and PPC (Fig. 6C, lower right panel). These results suggest that, in addition to the afferent input from the OB, VON receives inputs from various areas of the olfactory cortex, a part of the diagonal band, and the prefrontal cortex, as well as supporting the idea that the VON is a part of the olfactory cortex.

### Rabies virus-mediated trans-synaptic retrograde labelling from the LH to the VON

CTB labelling could not discriminate whether the labelled cells had synaptic connections to the postsynaptic neurons or whether their axon fibres were passing by in the injection site^28^. To address whether the VON neurons formed synaptic contacts in the LH, we used trans-synaptic retrograde labelling from the LH with the modified rabies virus as shown in Fig. 6. We injected Cre-encoding AAV and Cre-dependent AAVs encoding TVA-mCherry and rabies glycoprotein into the LH, followed by injection of the modified rabies virus (SAD-dG-EGFP+EnvA) into the LH^22^-^25^ (Fig. 7A). Because trans-synaptic spread of the rabies virus is mediated by the rabies glycoprotein^25^, we compared efficacy of retrograde labelling with and without rabies G expression in the LH. EGFP(+) cells were observed in the VON when we concomitantly injected rabies G-coding AAV into the LH (Fig. 7B, upper panels). In contrast, EGFP(+) cells were never observed in the VON in the absence of rabies G-coding AAV in the LH (Fig. 7B, lower panels). TVA-mCherry(+) and EGFP(+) cells in the LH were observed in cases with and without rabies G-coding AAV (data not shown). The results of rabies glycoprotein-dependent retrograde spread of rabies virus supports the idea that neurons in the VON form synaptic contacts in the LH.

**Figure 7.**
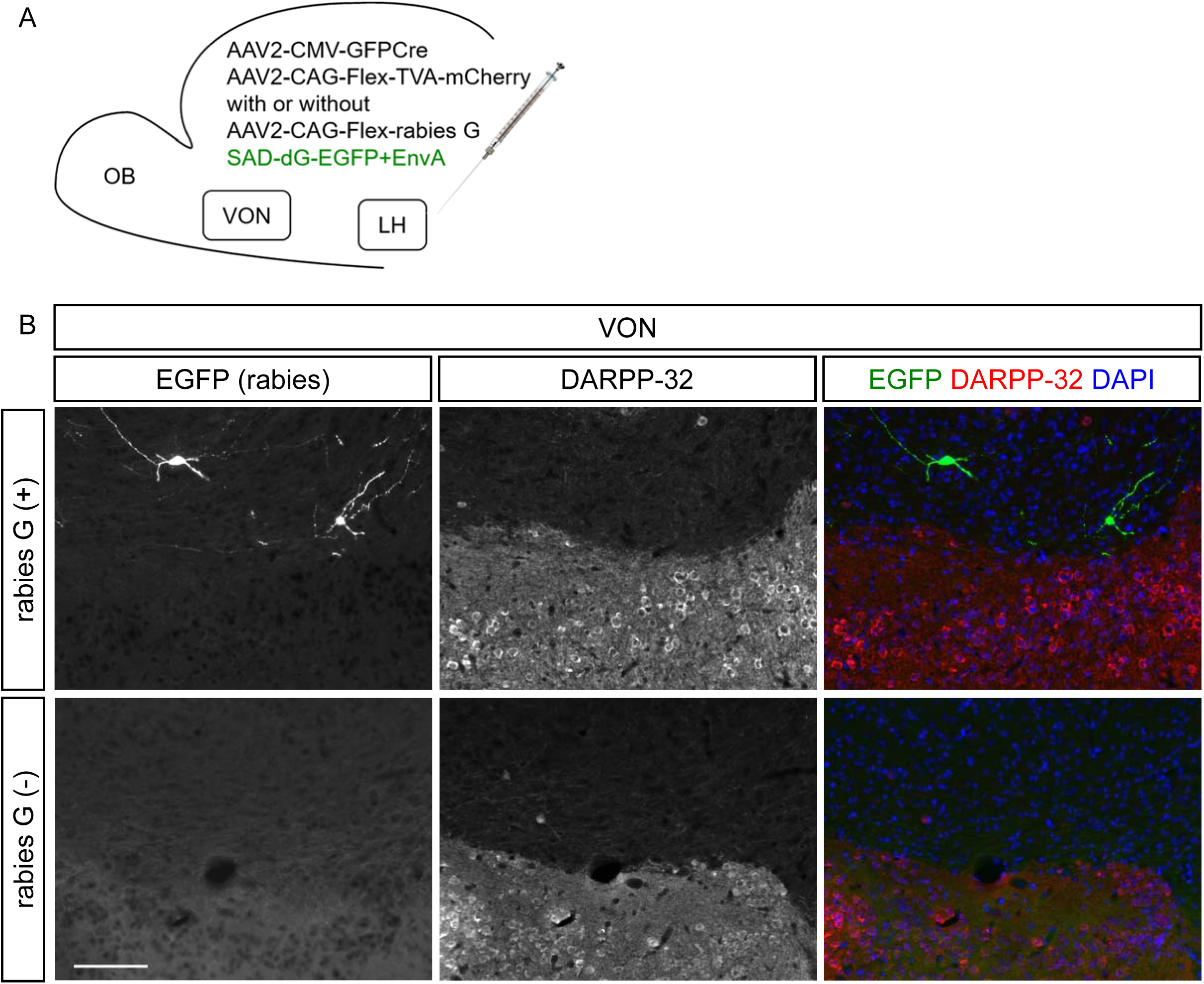
Trans-synaptic retrograde spread of rabies virus from the LH to the VON. (A) Schema of virus-mediated retrograde tracing. We first injected a mixture of adeno-associated viruses (AAVs) encoding CMV-GFPCre and CAG-Flex-TVA-mCherry either with or without CAG-Flex-rabies G into the lateral hypothalamus (LH). Two weeks later, SAD-dG-EGFP-EnvA was injected into the LH and we observed the distribution of EGFP-labelled presynaptic cells in the ventral olfactory nucleus (VON). (B) EGFP-labelled presynaptic neurons (left, green) were observed in the VON when rabies G-encoding AAV was concomitantly injected (upper panels). Immunostaining for DARPP-32 (middle, red) was used to discriminate the VON from the olfactory tubercle (OT) and nucleus accumbens (NAc). Scale bar: 100 μm.

## Discussion

In this study, we examined the neural pathways from the olfactory cortex to the LH in mice by retrograde tracing from the LH. We observed that a group of retrogradely labelled cells was clustered in a postero-ventral region of the olfactory peduncle. Our neuroanatomical and histochemical analyses revealed that this region predominantly comprises GABAergic neurons, thereby distinguishing it from the AON and VTT in which the principal neurons are glutamatergic. Furthermore, this region receives synaptic inputs from the M/TCs. These results suggest a novel population of GABAergic neurons in the olfactory cortex that project to the LH.

In accordance with the previous report by Price et al. in rats^11^, we identified an equivalent cluster of retrogradely-labelled cells in a posterior part of the olfactory peduncle in mice surrounded by the AON, VTT, APC, and OT (Figs. 1B and 3). Although it has been thought that this cluster of neurons belongs to the AON or VTT whose principal neurons are glutamatergic, our histochemical analyses showed that the majority of the neurons in this region were GABAergic, indicating that the retrogradely-labelled GABAergic projection neurons are distinct from glutamatergic projection neurons in the AON and VTT. We therefore temporarily named this region with GABAergic projection neurons the ventral olfactory nucleus (VON) which was located just beneath the AON. Then we examined neuroanatomical features of the VON.

It might be possible that the VON neurons are anteriorly displaced GABAergic medium spiny neurons of the OT and NAc because the location of the VON is just above the anterior tip of the OT and anterior to the NAc (Fig. 3). To examine this possibility, we performed immunostaining for DARPP-32 and found that the strongly immunoreactive OT and NAc neurons are clearly distinguishable from the less immunoreactive VON neurons (Figs. 3 and 4).

Fluorescent labelling confirmed synaptic contacts between M/TCs in the OB and VON neurons (Figs. 4 and 5). The dendrites of VON neurons projected in an antero-ventral direction toward the cortical surface and extended through the layer Ia of the olfactory cortex, where the dendrites were contacted by axons of M/TCs (Fig. 4). Dendritic spines of VON neurons were apposed to immunoreactive VGluT1 elements in the axons of M/TCs (Fig. 5). In addition to morphological analysis using retrograde AAV encoding EGFP and tdTomato-expressing transgenic mice, we used a modified rabies virus to demonstrate that M/TC axons form synaptic contacts with the VON neurons (Fig. 6). Taking advantage of the neuronal pathway-specific infection of the rabies virus combining retrograde Cre viral vectors and Cre-dependent expression of viral receptors and glycoproteins^26^, we successfully infected VON neurons with the modified rabies virus. We observed that M/TCs in the OB were labelled by this retrograde trans-synaptic method. These results confirmed that the VON is a part of the olfactory cortex receiving axonal inputs from M/TCs.

The disynaptic pathway from the OB to the LH via the VON seems to be one of shortcut pathways conveying odorant information to the LH. Unfortunately, the efficacy of our rabies virus mediated-labelling of presynaptic M/TCs was insufficient to address whether topographic axonal projections from the OB to the VON existed, as for other subregions of the olfactory cortex^29^. It remains to be addressed what odorant information the VON conveys to the LH via GABAergic outputs. In addition to the OB, the VON receives inputs from other regions of the olfactory cortex including the AON, APC, OT, and PPC, as well as modulatory inputs from the horizontal limb of the diagonal band, and top-down inputs from the prefrontal cortex (Fig. 6C). This organization suggests that VON neurons integrate these inputs and send GABAergic inhibitory output to the LH. Further studies should address the roles of each input to the VON neurons and how the VON neurons integrates these inputs.

Rabies virus-mediated trans-synaptic labelling from the LH suggested that VON neurons formed synaptic contacts in the LH (Fig. 7). The LH consists of a variety of neuronal subtypes including orexin neurons, MCH neurons, and GABAergic neurons, which have distinct roles in eating as well as sleep/wakefulness^12,13,30^. It was recently reported that medium spiny neurons in the NAc expressing dopamine receptor D1 have synaptic inputs to GABAergic neurons in the LH^31^. This NAc to LH pathway is involved in downregulation of feeding behaviour. Because our analysis did not identify the neuronal subtypes in the LH that receive GABAergic inputs from the VON, further neuroanatomical studies should address this point. In addition to anatomical analyses, functional manipulation by optogenetics and/or pharmacogenetics will help to reveal the roles of the VON. Our findings regarding the VON provide novel insight into the circuitry that may underpin odour-induced LH-related behaviours.

## Materials and Methods

### Animals

All experiments were conducted in accordance with the Guidelines for Animal Experimentation in Neuroscience of the Japan Neuroscience Society and were approved by the Experimental Animal Research Committee of University of Fukui and Doshisha University. C57BL/6J male mice were purchased from Japan SLC. Homozygote Pcdh21-nCre mice (C57BL/6Cr-Tg(Pcdh21-cre)BYoko RBRC02189, RIKEN BRC)^18^ and homozygote Ai14 mice (B6;129S6-Gt(ROSA)26Sortm14(CAG-tdTomato)Hze/J 007908, The Jackson Laboratory)^17^ were crossed. Male heterozygote mice for both genes were used for experiments in Figs. 4 and 5. All animals were individually housed after surgery with a 12/12 hour light/dark cycle. Food and water were available *ad libitum*.

### Virus preparation

For AAV vectors, AAVrg-CAG-GFP was a gift from Edward Boyden (Addgene viral prep #37825-AAVrg). AAV2-CMV-GFPCre, AAV2-CAG-Flex-TVA-mCherry, and AAV2-CAG-Flex-rabies G were packaged and concentrated to titres of 5.0 x 10^13^, 3.3 x 10^12^, and 1.3 x 10^12^ viral genomes/mL as previously reported^32^ using Addgene plasmids AAV-GFP/Cre (gift from Fred Gage, #49056^33^), CAG-Flex-TCB (gift from Liqun Luo, #48332^23^), and pAAV-CAG-FLEX-oG-WPRE-SV40pA (gift from Edward Callaway, #74292^22^), respectively.

For the rabies virus, we obtained EnvA-pseudotyped glycoprotein-deleted rabies virus encoding EGFP (SAD-dG-EGFP+EnvA) from Gene Transfer, Targeting and Therapeutics Facility of Salk Institute for Biological Studies at a titre of 2.9 x 10^7^ TU/mL.

For the lentivirus, NeuRet-Cre was packaged and concentrated to a titre of 1.1 x 10^12^ copies/mL as previously reported^32^.

### Stereotaxic surgery

The stereotaxic surgeries were performed on mice aged 10-16 weeks. Mice were anesthetized with a mixture of three anaesthetics (medetomidine, midazolam, and butorphanol)^34^, and then placed in a stereotaxic apparatus (Narishige, SR-5M). The skull above the targeted areas was thinned with a dental drill and removed carefully. Injections were conducted with a syringe pump (WPI, UltraMicroPump III) connected to a Hamilton syringe (Hamilton, RN-1701), mounted glass micropipette with a tip diameter of 50 μ m connected by an adaptor (Hamilton, 55750-01).

For Figs. 1-3, we unilaterally injected 150 nL of CTB conjugated with Alexa 555 into the left LH with the following coordinates: A/P, −1.2 mm and M/L, 1.2 mm from bregma; D/V, 4.8 mm from the brain surface. One week later, the mice were deeply anaesthetised and fixed as described below.

For Figs. 4 and 5, we unilaterally injected 300 nL of AAVrg-CAG-EGFP into the left LH of the double transgenic mice with the following coordinates: A/P, −1.2 mm and M/L, 1.2 mm from bregma; DV, 4.8 mm from the brain surface. Two weeks later, the mice were deeply anesthetized and fixed as described below.

For Fig. 6, we unilaterally injected 300 nL of NeuRet-Cre into the left LH and 300 nL of 1:1 mixture of two AAVs (AAV2-CAG-Flex-TVA-mCherry and AAV2-CAG-Flex-rabies G) into the left VON. For the LH, we used the following coordinates: A/P, −1.2 mm and M/L, 1.2 mm from bregma; DV, 4.8 mm from the brain surface. For the VON, we used the following coordinates: A/P, +2.0 mm and M/L, 0.8 mm from bregma; DV, 4.2 mm from the brain surface. Two weeks later, we unilaterally injected 300 nL of SAD-dG-EGFP+EnvA using the same VON coordinates. One week later, the mice were deeply anaesthetised and fixed as described below.

For Fig. 7, we unilaterally injected 300 nL of 1:1:1 mixture of three AAVs (AAV2-CMV-GFPCre, AAV2-CAG-Flex-TVA-mCherry, and AAV2-CAG-Flex-rabies G) for rabies G(+) mice or 300 nL of 1:1 mixture of two AAVs (AAV2-CMV-GFPCre and AAV2-CAG-Flex-TVA-mCherry) for rabies G(-) mice into the left LH with the following coordinates: A/P, −1.2 mm and M/L, 1.2 mm from bregma; D/V, 4.8 mm from the brain surface. Two weeks later, we unilaterally injected 300 nL of SAD-dG-EGFP+EnvA using the same LH coordinates. One week later, the mice were deeply anaesthetised and fixed as described below.

### Sample preparation for histochemistry

Mice were deeply anaesthetised by intraperitoneal injection of sodium pentobarbital. They were transcardially perfused with phosphate-buffered saline (PBS) followed by 4% paraformaldehyde (PFA). The brains were removed from the skull, immersed in 4% PFA in 0.1 M phosphate buffer (PB) overnight, and then transferred to 30% sucrose in 0.1 M PB. The brains were then embedded in O.C.T. compound (Sakura Finetechnical), frozen at −80°C, and sliced into coronal sections with a thickness of 20 μm with a cryotome. Sections were rinsed in PBS and 0.1 M PB, mounted on glass slides (Matsunami, CREST) using a paint brush, dried overnight in a vacuum desiccator, and then stored at 4°C until histochemistry.

### Histochemistry

We performed immunostaining for orexin (Fig. 1A, middle), MCH (Fig. 1A, right), DARPP-32 (Figs. 3, 4, and 7), EGFP (Figs. 4 and 5), and VGluT1 (Fig. 5) as follows. The dried sections were rehydrated in PBS, permeabilised in TNT (0.1 M Tris-HCl; pH, 7.5; 0.15 M NaCl; 0.1% Tween 20), and blocked with 10% normal donkey serum diluted in TNT. Then, the sections were incubated with the following primary antibodies overnight at 4°C: goat anti-orexin polyclonal antibody (1:400; Santa Cruz sc-8070); rabbit anti-MCH polyclonal antibody (1:400, Sigma M8440); rabbit anti-DARPP-32 monoclonal antibody (1:400; Abcam ab40801); rat anti-EGFP monoclonal antibody (1:1000, Nacalai Tesque 04404-84); guinea pig anti-VGluT1 polyclonal antibody (1:500, Merck AB5905). After three washes in TNT, appropriate fluorescent dye-conjugated secondary antibodies (1:400; Jackson ImmunoResearch) were incubated for 2 hours at room temperature. After three washes in TNT, the sections were then counterstained with DAPI diluted in PBS (2 µg/mL) for 5 min. After washing in PBS, the sections were mounted in PermaFluor (Thermo Fisher Scientific).

For Fig. 2A, we performed *in situ* hybridization for VGluT1, VGluT2, VGluT3, and GAD65/67 mRNA as follows. Digoxigenin (DIG)-labelled RNA probes were made using an *in vitro* transcription kit (Roche) according to the manufacturer’s protocol with plasmids kindly provided by Drs. Katsuhiko Ono and Yuchio Yanagawa^35^-^37^. The dried sections were fixed in 4% PFA, digested with Proteinase K (10 μg/mL) for 30 min, and post-fixed in 4% PFA. After prehybridization, the sections were incubated overnight at 65°C with DIG-labelled RNA probes. After stringent washing, the sections were blocked with 10% normal sheep serum, 1% bovine serum albumin (BSA), and 0.1% Triton X-100 in PBS. Subsequently, the sections were incubated overnight at 4°C with alkaline phosphatase-conjugated anti-DIG antibody (1:1000; Roche). The sections were washed in TNT, followed by alkaline phosphatase buffer (100 mM NaCl; 100 mM Tris-HCl; pH, 9.5; 50 mM MgCl2; 0.1% Tween 20; 5 mM levamisole). The sections were treated overnight with NBT/BCIP (Roche) mixture at room temperature in a dark room for colour development. Then, they were rinsed in PBS and mounted in PermaFluor (Thermo Fisher Scientific).

For Fig. 2B, we performed double fluorescent labelling of immunoreactivity for CTB and mRNA for VGluT1 or GAD65/67 as follows. Fluorescein-labelled RNA probes were prepared as described above. Hybridization and washing were performed as described above, except that fluorescein-labelled probes were used for hybridization. After blocking in 1% blocking buffer (11096176001, Roche) for 1 h, the fluorescent-labelled probes were detected. The sections were incubated with an anti-fluorescein antibody conjugated with horseradish peroxidase (1:500; Perkin-Elmer) for 1 h at room temperature. After three 10-min washes in TNT, the sections were treated with diluted (1:100) TSA-Plus dinitrophenol (DNP) reagents for 5 min according to the manufacturer’s instructions (Perkin-Elmer), and the FLU signals were converted to DNP signals. To amplify the DNP signals, the sections were washed in TNT three times for 10 min each, incubated with an anti-DNP antibody conjugated with horseradish peroxidase (1:500; Perkin-Elmer) for 1 h at room temperature, and treated again with diluted TSA-Plus DNP reagents (1:100) for 5 min. Subsequently, the sections were incubated overnight with an anti-DNP antibody conjugated with Alexa 488 (1:500; Molecular Probes) in 1% blocking buffer at 4°C for fluorescence detection of DNP signals. At this point, a goat anti-CTB antibody (1:500; List Biological Laboratories #703) was added to the incubation mixture for detection of CTB. The sections were washed three times in TNT and incubated with a Cy3-conjugated secondary antibody (1:400; Jackson ImmunoResearch Labs) for 2 h. After three washes in TNT, the sections were then counterstained with DAPI diluted in PBS (2 µg/mL) for 5 min. After washing in PBS, the sections were mounted in PermaFluor (Thermo Fisher Scientific).

For Figs. 1A (left panel), 1B, 2A (two left panels), and 6, the dried sections were rehydrated in PBS and counterstained with DAPI diluted in PBS (2 µg/mL) for 5 min. After washing in PBS, the sections were mounted in PermaFluor (Thermo Fisher Scientific).

### Microscopy

Sections were examined with a bright field virtual slide system (Hamamatsu Photonics, NanoZoomer), a fluorescent microscope (Olympus, BX51WI), and a confocal laser microscope (Olympus, FV1200).

## Availability of Materials and Data

In publication we make materials, data and associated protocols promptly available to readers without undue qualifications in material transfer agreements.

## Acknowledgements

We thank Drs. Edward Callaway, Fumitaka Osakada, and Akihiro Yamanaka for providing viral vectors and valuable discussion, Drs. Katsuhiko Ono and Yuchio Yanagawa for plasmids for RNA probes, Eri Murai, Noriko Funabashi, and members of the Fukazawa laboratory and Life Science Research Laboratory at University of Fukui for technical assistance. K.Murata was supported by the Takeda Science Foundation, Cosmetology Research Foundation, and JSPS KAKENHI Grant Numbers 16K18377, 16H01671, 17KK0190, 18H05005. Y.F. was supported by JSPS KAKENHI Grant Numbers 15H01282, 16H04662. H.M. was supported by the Takeda Science Foundation, Urakami Foundation for Food and Food Culture Promotion and JSPS KAKENHI Grant Numbers 16K14557, 16H02061.

## Author Contributions

K.Murata and H.M. designed research, performed experiments, analysed data, and wrote the paper. T.K. performed experiments. Y.F., Kenta.K., Kazuto K., K.Miyamichi, H.O., H.B., Y.S., M.Y., and K. Mori contributed to reagents and commented on the manuscript.

